# The quest continues: Human CD4^+^ CD16^+^ CD56^+^ ‘exTreg’ resemble NKT cells instead

**DOI:** 10.1101/2025.08.05.667656

**Authors:** Jeevan Mutha, Meghana Konga, Maximilian Sprang, Johannes U Mayer, Susan M Schlenner

## Abstract

Regulatory T cells are essential for immune tolerance, but their loss of function under inflammatory conditions in murine models signify a risk factor for Treg-based therapies. Recently CD4^+^ CD56^+^ CD16^+^ T cells were suggested to resemble such ex-Treg in human PBMC. Here, we re-evaluate the identity of the CD4^+^ CD56^+^ CD16^+^ population at a phenotypic and transcriptomic level using multiparametric flow cytometry on human PBMC and CITE-seq analysis to demonstrate that the CD4^+^ CD56^+^ CD16^+^ cells mostly constitute NKT cells instead. Further, we evaluated the stability of human Treg under lineage-challenging conditions and observe robust lineage stability in vitro. Finally, we also explore the potential of Tr17 induction using TGF-β and IL-6, a possible therapeutic strategy for Treg ex vivo expansion-based therapies. Together, we conclude that human exTreg remain to be described and instead human Treg present as remarkably stable, further promoting Treg-based adoptive transfer therapies.

## Introduction

Regulatory T cells (Treg) play a prominent role in maintaining immune tolerance and preventing autoimmunity, primarily through the suppression of effector T cell responses^1^. Their suppressive functions are multifaceted and are mediated via both cell contact-dependent mechanisms, involving molecules such as CTLA-4 and CD39^2,3^, and the secretion of soluble factors, including immunosuppressive cytokines like TGF-β, IL-10, and IL-35^4–6^. Additionally, Treg can suppress pro-inflammatory T cell proliferation through IL-2 sequestration^7^. Deficiencies in Treg functional genes, notably, disruptions in the *FOXP3* gene cause the Immune Dysregulation Polyendocrinopathy Enteropathy X-linked (IPEX) syndrome, a severe autoimmune condition^1^. Consequently, Treg-based and Treg-targeted therapies are increasingly being researched, with several ongoing trials to treat autoimmune conditions^8–10^.

Foxp3 is the master transcription factor in Treg which mainly acts as a transcriptional repressor of pro-inflammatory genes such as *Il2*, *Ifng*, *Zap70*^11^. Stable expression of Foxp3 is necessary to maintain the suppressive capabilities of mature Treg in the periphery^12^. IL-2/CD25 signalling through pSTAT5 maintains this stable expression by inhibiting the re-methylation of the *Foxp3* locus^11,13^. Despite the requirement of stable Foxp3 expression, under inflammatory conditions, Treg can upregulate lineage-defining transcription factors and chemokine receptors characteristic of T_H_ cell subsets, such as T-bet (T_H_1), GATA3 (T_H_2), or RORγt (T_H_17), while retaining Foxp3 expression. This functional adaptation is driven by the same cytokine factors that drive specific T_H_-subset differentiation during inflammation and is required for T_H_-subset-specific immunosuppression^14–17^. The Foxp3 transcriptional circuit and the proinflammatory transcriptional profile driven by the T_H_-specific transcription factors co-exist in competition and in a delicate balance in T_H_-like Treg, hence allowing the T_H_-like Treg to carry out immunosuppressive function without contributing to inflammation^13^.

However, this plastic nature of Treg can result in loss of Foxp3 expression and acquisition of a pro-inflammatory phenotype due to imbalances of Treg lineage-maintenance and pro-inflammatory signals^13,18,19^. This has been best described for the Treg-T_H_17 axis, where Tr17 (RORγt^+^ Treg) develop to control T_H_17-driven inflammation with key cytokines being TGF-β, IL-6, and IL-2^13^. While TGF-β together with IL-2 promotes their differentiation into Foxp3⁺ Treg, TGF-β with IL-6 favours T_H_17 differentiation by inducing RORγt and IL-17. This is controlled by the opposing actions of STAT5 and STAT3, downstream of IL-2 and IL-6 respectively, which bind to overlapping DNA sites with largely opposing functions. The balance between STAT5 and STAT3 signalling determines the functional phenotype along the Treg-T_H_17 axis^20,21^ and an imbalance favouring IL-6-mediated STAT3 signalling can result in formation of T_H_17-like exTreg cells in mice^22^. Mechanistically this is mediated through the IL-6-driven phosphorylation of STAT3, increased expression of the transcriptional regulator Id2, which negatively regulates Foxp3 expression and can re-methylate the *Foxp3* locus through an increase in expression of DNMTs^22–26^.

For Treg-based therapies which require the purification, potential *ex vivo* manipulation, and extensive expansion of Treg and their reintroduction into patients, warranting Treg stability is of central importance^27,28^. Even more so, as Treg derived from thymus are reactive to self-antigens^29^, and loss of FOXP3 expression and a subsequent adoption of a pro-inflammatory features can potentially exacerbate conditions that Treg-based therapies aim to treat.

Most of the studies on exTreg have been conducted in mice, however, the existence of exTreg in humans remains contentious. Freuchet et al. described the phenotype of human exTreg as CD56^+^ CD16^+^ cytotoxic T cells, also circulating in the blood of healthy human PBMC donors. The authors concluded this from correlating the transcriptomic signature of exTreg in atherosclerotic *Apoe^-/-^* mice with Treg fate mapping (FoxP3^eGFP−Cre-ERT2^ ROSA26^fl-stop-fl-tdTomato^) to a similar T cell population in a published CITE-seq dataset from coronary artery disease (CAD) patients. These cells were shown to express granzyme B, perforin, and T_H_-like cytokines and expressed canonical NK-related genes with T cell functionality^30^. The authors carefully disproved that the discovered cell population was distinct from NK cells. However, the described exTreg associated features also align with Natural killer (NK) T lineage, which are characterized by expression of CD3 and CD56, can exert cytotoxic effects in a CD1d-dependent (type I NKT, also called invariant NKT, and type II NKT) and CD1d-independent and MHC-unrestricted manner (NKT-like), and exist in CD4^+^, CD8^+^ and CD4/CD8 double-negative populations^31–33^.

To clarify to what extent the proposed CD4^+^ CD56^+^ CD16^+^ exTreg population does align with features of NKT cells in healthy human PBMC we delineated the population based on the NKT lineage markers Promyelocytic Leukaemia Zinc Finger (PLZF), CD161 and activation phenotype markers. Further, we analysed published human exTreg transcriptomic and NKT signatures in CD4^+^ CD56^+^ CD16^+^ cells using various published CITE-seq datasets and strikingly observed that the proposed exTreg signature fails to characterize a specific cluster within CD56^+^ CD16^+^ cells nor differentiates from the NKT cell profile. Furthermore, cultured human Treg proved to be highly stable – even under lineage fate-pressurizing conditions – suggesting that human Treg may be more stable than their murine counterparts and that exTreg do not represent a sizable population in human PBMC.

## Results

### Human CD3^+^ CD4^+^ CD56^+^ CD16^+^ represent a heterogenous population, resembling NKT

Human NKT cell development involves PLZF and CD161 (NK1.1), and the expression of these lineage-defining markers is maintained in the periphery^34^. NKT cells show remarkable functional diversity in terms of their ability to produce T_H_-like cytokines and are also capable of producing the immunosuppressive cytokine IL-10^34,35^. To understand the phenotype of human CD3^+^ CD4^+^ CD56^+^ CD16^+^ cells and determine if they consist both exTreg and NKT cell populations, we developed an antibody panel including the lineage markers PLZF, CD161 (NK, NKT, unconventional T cell lineages), CRTH2 (T_H_2); activation markers CD27, CD127, CD25, CD45RO, CCR7, as well as inhibitory/exhaustion markers PD-1, TIM-3 and TIGIT. Employing FlowSOM clustering, we analysed the phenotypes present in the CD3^+^ CD4^+^ CD56^+^ CD16^+^ population **(Fig. 1A, B)**. This analysis identified four clusters, with three clusters displaying high PLZF expression while differing in their activation status and one PLZF^neg^ expression cluster. PLZF expression levels in CD3^+^ CD4^+^ CD56^+^ CD16^+^ cells and distinctly gated CD3^+^ CD56^+^ NKT cells was comparable, while CD3^+^ CD4^+^ CD56^-^ CD16^-^ conventional T cells expressed low levels or PLZF **(Fig. 1C)**. CD3^+^ CD56^+^ PLZF^+^ population have been shown to have high granularity^31^ and indeed PLZF expression in the CD3^+^ CD4^+^ CD56^+^ CD16^+^ population correlated with high granularity **(Fig. 1D)**. In PBMC from healthy individuals the majority of the CD3^+^ CD4^+^ CD56^+^ CD16^+^ population expressed PLZF (61.85% ± 24.05%; mean ± SD, n=5) and most PLZF-expressing cells displayed a naïve CD45RO^-^ CCR7^+^ phenotype (35.84% ± 16.15%; mean ± SD, n=5), while 22.54% ± 9.05% (mean ± SD, n=5) presented an activated/memory phenotype (CD45RO^+^) **(Fig. 1E)**.

**Figure 1.**
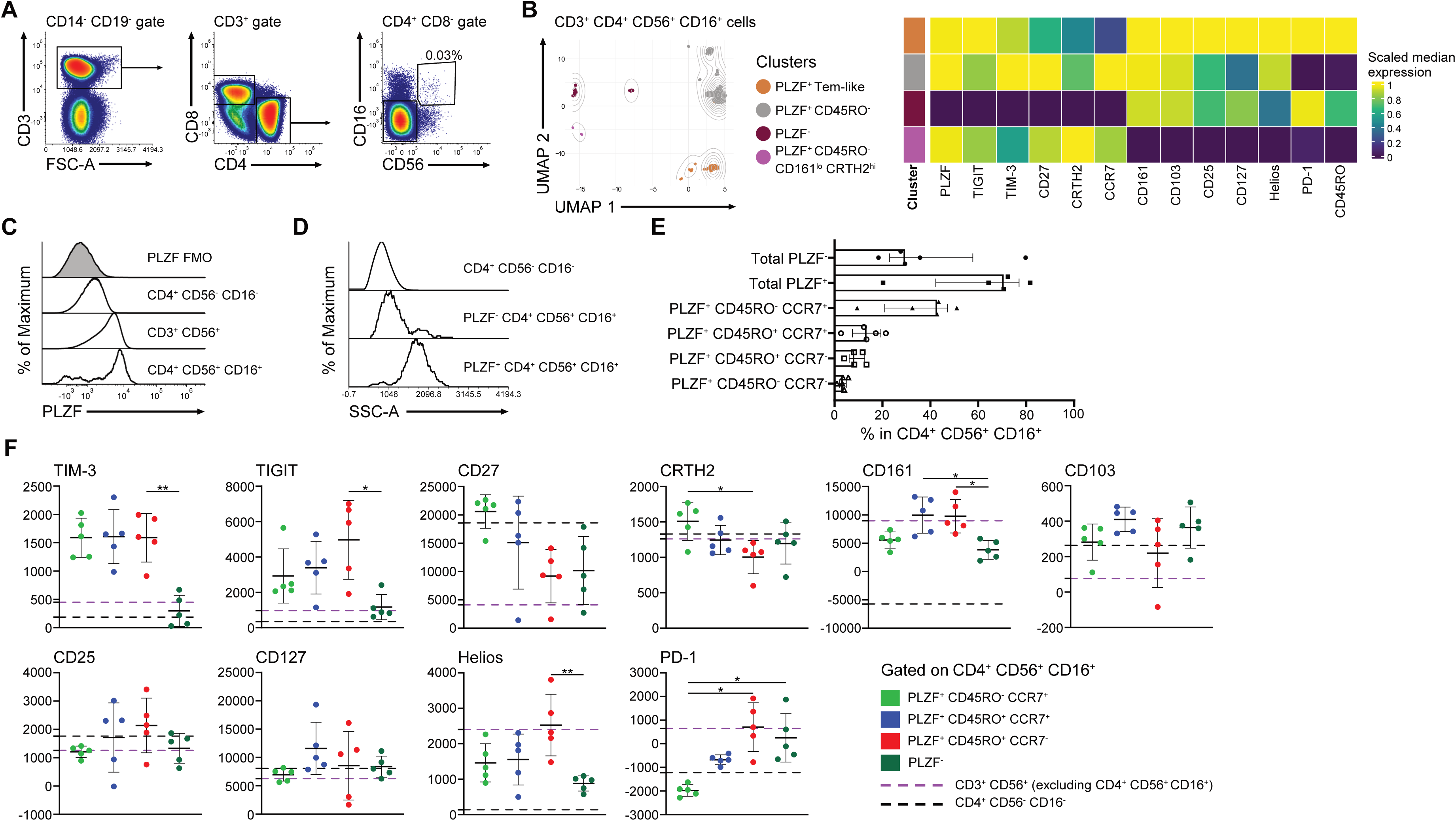
Human CD3^+^ CD4^+^ CD56^+^ CD16^+^ cells express PLZF and resemble activated and naïve NKT cells. **(A)** Representative flow cytometry gating strategy for identification of CD3^+^ CD4^+^ CD56^+^ CD16^+^ cells from healthy human PBMC. Refer to Suppl. Fig 1A for the pre-gating strategy. **(B)** UMAP embedding of FlowSOM clustering of CD3^+^ CD4^+^ CD56^+^ CD16^+^ cells (left) and scaled heatmap expression of phenotyping markers in the respective clusters (right), pooled from 5 individual samples. **(C)** Representative histogram overlay of PLZF expression in conventional T cells (CD3^+^ CD4^+^ CD56^-^ CD16^-^), CD3^+^ CD4^+^ CD56^+^ CD16^+^ cells and CD3^+^ CD56^+^ NK cells (excluding CD4^+^ CD56^+^ CD16^+^ cells). **(D)** Representative histograms of granularity (SSC-A) within PLZF^+^ and PLZF^-^ CD3^+^ CD4^+^ CD56^+^ CD16^+^ cells and conventional T cells. **(E)** Bar graph proportion of PLZF, CD45RO, and CCR7 expressing cell populations within the CD3^+^ CD4^+^ CD56^+^ CD16^+^ cells (median ± IQR, n=5). **(F)** Scatter plots depicting MFI of immunophenotyping markers in the PLZF^-^ and PLZF^+^ subpopulations of CD3^+^ CD4^+^ CD56^+^ CD16^+^ cells (mean ± SD, n=5). Median expression levels by conventional CD4^+^ CD56^-^ CD16^-^ T cells and CD3^+^ CD56^+^ NK cells (excluding CD4^+^ CD56^+^ CD16^+^) are depicted as lines. Statistical analysis for comparison of the three groups was performed using the Friedman test with post-hoc Dunn’s multiple comparisons. [*p<0.05, **p<0.01; only significant comparisons are indicated.]

Further analysis of PLZF^+^ CD45RO^-^ CCR7^+^ naïve-like, PLZF^+^ CD45RO^+^ CCR7^+^ Tcm-like, PLZF^+^ CD45RO^+^ CCR7^-^ Tem-like, and PLZF^-^ populations within the CD3^+^ CD4^+^ CD56^+^ CD16^+^ cells revealed that TIM-3 has a higher baseline expression in PLZF^+^ populations compared to PLZF^-^ populations, which has been previously attributed as a feature of the NK lineage^36^. We also observe a lower expression of the NK, NKT lineage marker CD161, TIGIT and HELIOS in the PLZF^-^ population compared to PLZF^+^ populations. Expression of PD-1 tended to be higher in the PLZF^+^ CD45RO^+^ CCR7^-^ Tem-like cells and PLZF^-^ cells compared to PLZF^+^ CD45RO^+^ CCR7^+^ Tcm-like and CD45RO^-^ CCR7^+^ naïve-like cells. **(Fig. 1F, Suppl. Fig. 1B)**.

In murine studies, exTreg are characterized by a loss of FOXP3 expression^22^, while both human and mouse NKT cells can express FOXP3 through mechanisms including TCR signalling, stimulation with TGF-β, rapamycin, or IL-10^37–39^. We therefore analysed FOXP3 and CD25 co-expression in the CD3^+^ CD4^+^ CD56^+^ CD16^+^ population and observed that FOXP3^+^ CD25^+^ cells are present at frequencies comparable to that of Treg among CD3^+^ CD4^+^ Tconv cells **(Suppl. Fig. 1C)**. Additionally, in both FOXP3^+^ CD25^+^ populations, the majority of cells expressed low levels of CD127, while FOXP3^+^ CD25^+^ cells within the CD3^+^ CD4^+^ CD56^+^ CD16^+^ population expressed high PLZF and CD161 **(Suppl. Fig. 1D)**.

Overall, our results show that in PBMC of healthy individuals CD3^+^ CD4^+^ CD56^+^ CD16^+^ cells are dominated by NKT cells, identified by their lineage markers PLZF and CD161, which can be subdivided into activated/memory and naïve phenotypes and a population of FOXP3^+^ CD25^+^ cells. This heterogeneity suggests that an exTreg population, if indeed exists, requires further elaboration for accurate identification within CD3^+^ CD4^+^ CD56^+^ CD16^+^ cells and can likely not be identified by surface markers alone.

### Cells expressing the transcriptomic ‘exTreg signature’ *CST7*, *NKG7*, *GZMA*, *PRF1*, *TBX21*, *CCL4* also express NKT signature genes

In order to determine if the trancriptomic exTreg signature published by Freuchet et al^30^ highlights unique cells distinct from NKT we analysed the Coronary Artery Disease (CAD) patient CITE-seq dataset originally used to determine the ‘exTreg phenotype’^30^. We also analysed four additional PBMC datasets with suitable CITE-seq parameters, including one dataset with healthy volunteers enrolled in a HIV vaccine trial (VAC), and three datasets involving patients with immune-related disease conditions (type-1 diabetes [T1D], psoriatic arthritis [PsA], *ARPC5* mutation [ARPC5mut] datasets)^40–43^. CD3^+^ CD4^+^ CD56^+^ CD16^+^ cells were selected based on the respective antibody-derived tags (ADTs) **(Fig. 2A)** and after an unbiased clustering analysis we identified two different cell clusters in CAD and HD – vaccine trial datasets, and three different cell clusters in T1D, PsA and ARPC5mut datasets (**Fig. 2B)**. Furthermore, ‘exTreg transcriptomic signature’ expression (*CST7*, *NKG7*, *GZMA*, *PRF1*, *TBX21*, *CCL4*) highly correlated (p<2.2e-16) with the expression of NKT signature genes (*NKG7*, *FGFBP2*, *GZMK*, *GZMB*, *CDC42*, *GZMH*, *KLRD1*, *KLRB1*, *LEF1*, *SELL*, *PRF1*, *TNFRSF4*, *CCL3L1*), previously published by Zhou et al^44^, in all the analysed CITE-seq datasets with correlation coefficients of R=0.8 (CAD), R=0.93 (VAC), R=0.61 (T1D), R=0.7 (PsA) and R=0.7 (ARPC5mut) datasets **(Fig. 2C, D)**. Gene expression analysis revealed that the two population clusters of the CAD dataset differentially express *TRAC* and *TRDC*, which encode the δ and α chains of TCR respectively, suggesting that a portion of the CD3^+^ CD4^+^ CD56^+^ CD16^+^ cells also contain γδ T cells **(Fig 2E)**, while cells highly expressing the exTreg transcriptomic signature tended to express lower levels of *TRAC* **(Fig. 2D, E)**. In the ARPC5mut dataset, the CITE-seq panel also included an antibody targeting invariant Vα24-Jα18 TCR chain, a marker of iNKT cells. Analysis of Vα24-Jα18 expression indicated that iNKT cells are highly enriched in the CD3^+^ CD4^+^ CD56^+^ CD16^+^ population, making up 71.94% of cells **(Fig. 2F, G)**, verifying that NKT are a major population of CD3^+^ CD4^+^ CD56^+^ CD16^+^ PBMC cells.

**Figure 2.**
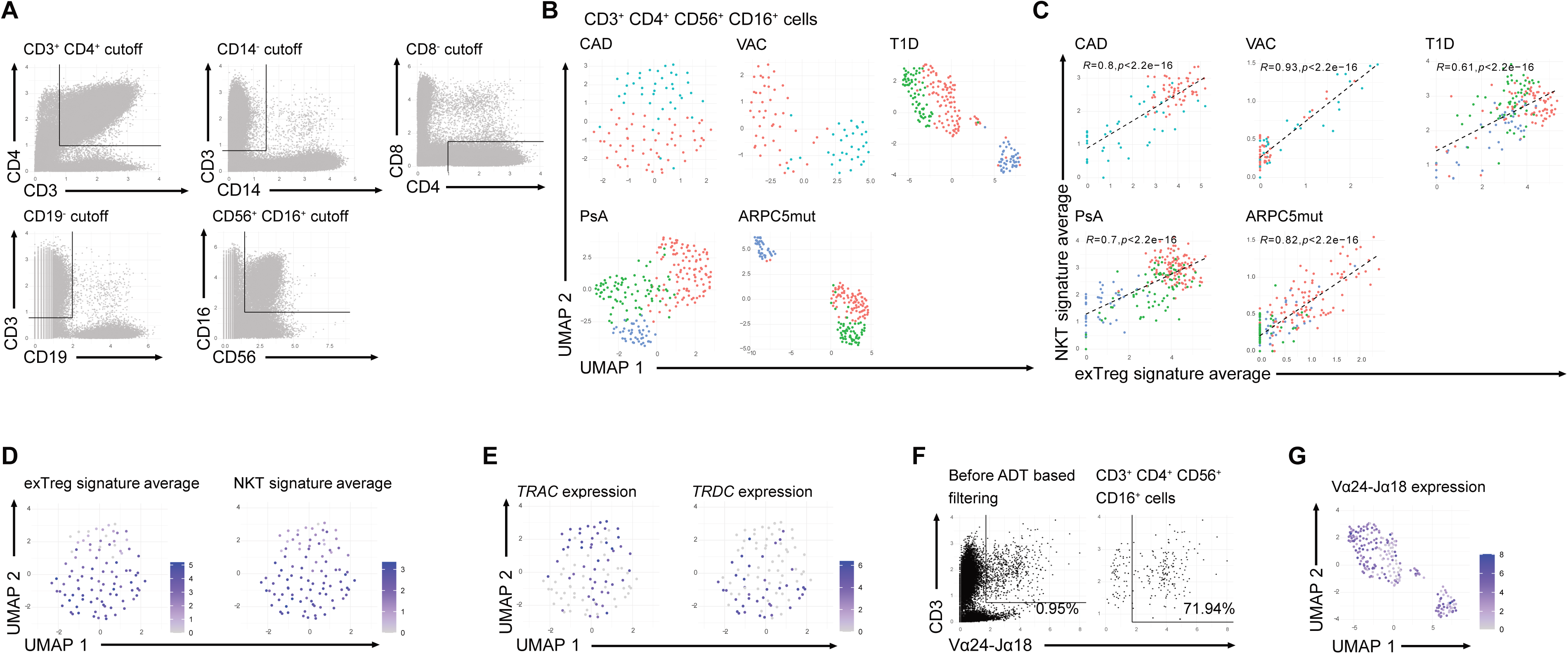
Analysis of exTreg and NKT transcriptomic signatures on CD3^+^ CD4^+^ CD56^+^ CD16^+^ cells from CITE-seq datasets. **(A)** Representative scatter plots displaying our ADT (antibody-derived tags) cut-off strategy for selecting CD3^+^ CD4^+^ CD56^+^ CD16^+^ CD14^-^ CD19^-^ CD8^-^ cells from CITE-seq datasets. **(B)** UMAP plots depicting clusters of CD3^+^ CD4^+^ CD56^+^ CD16^+^ cells from five CITE-seq datasets. CAD – Coronoary artery disease patients^30^, VAC – Healthy volunteers from HIV Vaccine trial^40^, T1D – Type-1 diabetes patients^41^, PsA – Psoriactic Arthritis patients^42^, ARPC5mut – Actin-related protein 2/3 complex subunit 5 mutation patients^43^. **(C)** Regression plots depicting the correlation between average expression of the published transcriptomic exTreg^30^ and NKT^44^ signatures in CD3^+^ CD4^+^ CD56^+^ CD16^+^ cells for all five CITE-seq datasets. Statistical analysis was performed using the pearson correlation method. **(D)** Average expression of exTreg and NKT transcriptomic signature genes overlaid on the UMAP plot of CD3^+^ CD4^+^ CD56^+^ CD16^+^ cells from the CAD CITE-seq dataset. **(E)** Expression levels of *TRAC* and *TRDC* overlaid on the UMAP plot of CD3^+^ CD4^+^ CD56^+^ CD16^+^ cells from the CAD CITE-seq dataset. **(F)** Scatter plots depicting ADT expression of CD3 and the iNKT-marker Vα24-Jα18 in the ARPC5mut CITE-seq dataset before and after filtering for CD3^+^ CD4^+^ CD56^+^ CD16^+^ cells. **(G)** Expression levels of Vα24-Jα18 overlaid on the UMAP plot of CD3^+^ CD4^+^ CD56^+^ CD16^+^ cells from the ARPC5mut CITE-seq dataset.

### IL-6Rα expression and suppressive marker profile of naïve and activated Treg subsets

Following our above conclusions that exTreg in human PBMC are not readily detectable, we next asked whether human Treg expansion cultures could be used to study FOXP3 instability *in vitro*. Treg instability studies in mice suggest that naïve Treg populations with lower Foxp3 expression are more susceptible to become exTreg compared to activated Treg with higher Foxp3 expression^45^. Furthermore, IL-6 has been proposed as a strongly antagonising molecule in regard to Foxp3 expression^21^. Therefore, we first evaluated whether the level of IL-6Rα expression differs between naïve (CD45RA^+^) and activated/memory (CD45RA^-^) human Treg subsets and correlates with the presence of other Treg hallmarks. Treg from rested PBMC of healthy donors were characterized for the expression of TIGIT, CTLA-4, GITR, CCR4, CD39, CD25 and FOXP3, proteins that are important for suppressive capabilities or migration, as well as markers that distinguish naïve cells and recent thymic emigrants (RTE; CD31, CD45RA) **(Fig. 3A)**. FlowSOM clustering identified four distinct Treg subpopulations based on these markers, one cluster describing naïve Treg and three clusters describing different features of activated/memory Treg and as expected, we observed lower expression of the Treg functional markers in CD45RA^+^ subset compared to the CD45RA^-^ subsets **(Fig. 3B-D)**. Targeted profiling of IL-6Rα expression in naïve and activated/memory Treg populations from rested PBMC revealed that both CD45RA-positive and -negative Treg populations expressed IL-6Rα **(Fig. 3E-F)**. Activated Treg uniformly expressed IL-6Rα at significantly higher levels than naïve Treg, indicating that IL-6Rα expression is a common feature among activated/memory Treg **(Fig. 3G)**. Furthermore, CD45RA^+^ Treg frequency declined with donor age, which aligns with previous studies describing a reduced naïve Treg pool in the elderly^46^**(Fig. 3G)**. These results postulate that IL-6Rα^high^ CD45RA^-^ maybe more susceptible to IL-6-driven fate challenge – leading to the induction of functional activation programs – upon loss of FOXP3 expression.

**Figure 3.**
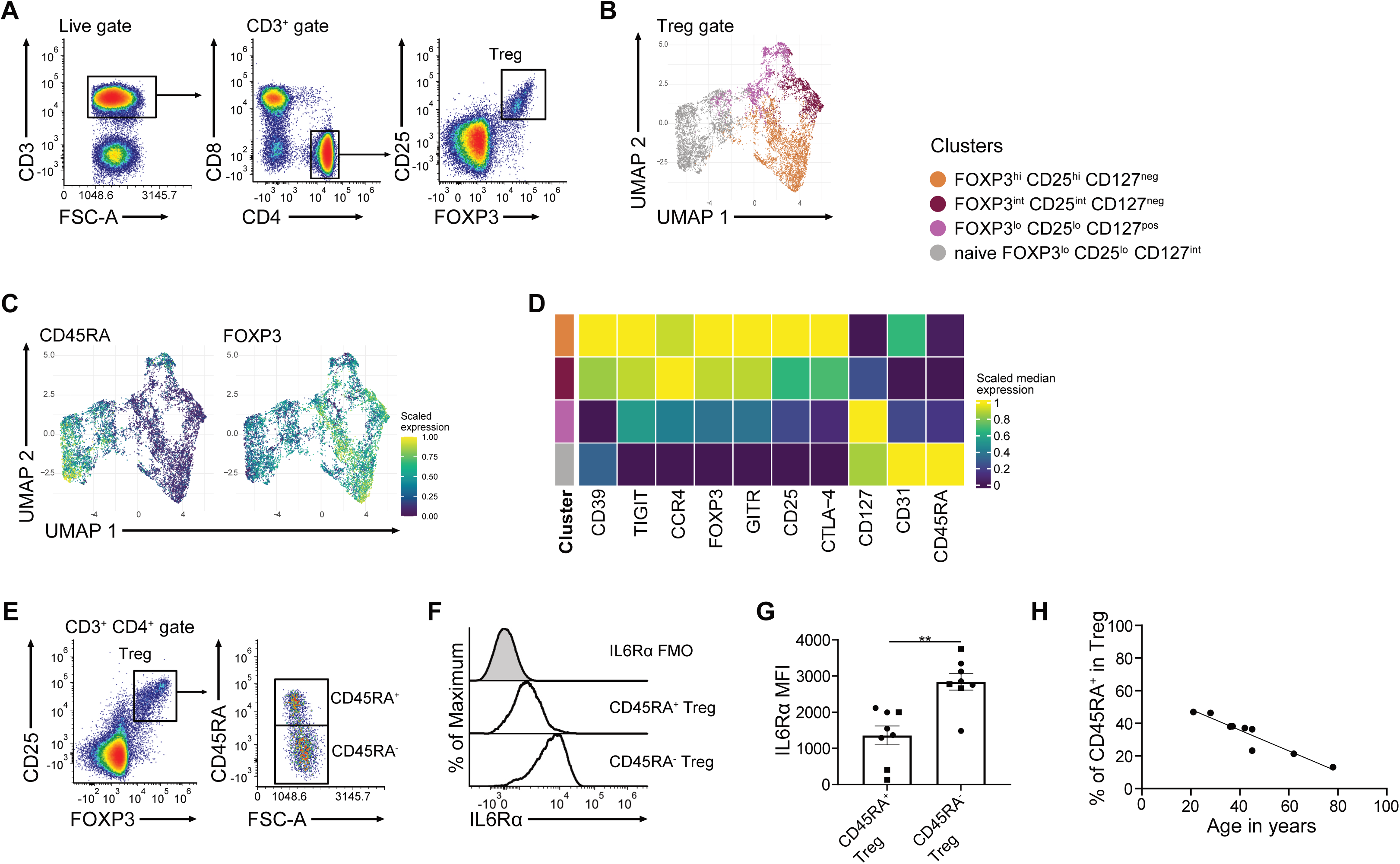
Profiling of FOXP3-low, intermediate and high expressing Treg and IL6Rα expression levels in PBMC of healthy donors. **(A)** Representative flow cytometry gating strategy for identification of FOXP3^+^ CD25^+^ Treg from human PBMC. n=8. **(B)** FlowSOM clustering overlaid on the UMAP plot of FOXP3^+^ CD25^+^ Treg, pooled from 8 PBMC samples. **(C)** Relative expression of CD45RA (left) and FOXP3 (right) overlaid on the UMAP plot of FOXP3^+^ CD25^+^ Treg, pooled from 8 PBMC samples. **(D)** Heatmap depicting relative median expression of Treg markers in each identified cluster of FOXP3-low, intermediate and high expressing Treg. **(E)** Representative gating strategy to identify naïve (CD45RA^+^) and activated/memory (CD45RA^-^) Treg subsets from human PBMC. **(F)** Representative histograms of IL6Rα expression in naïve and activated Treg. **(G)** IL6Rα MFI of naïve and activated Treg (mean ± SD, n=8) from two independent experiments, differentiated by the shape of data points. Wilcoxon matched-pairs signed rank test was performed for statistical analysis. **(H)** Linear regression plot of CD45RA^+^ cell frequencies in Treg against the age of 9 healthy PBMC donors. Statistical analysis was performed using the F-test. [**p<0.01; only significant comparisons are indicated.]

### Treg expansion cultures maintain lineage stability upon stability challenge with IL-6, even under limited IL-2 concentrations

Given the differential expression intensity of IL-6Rα on naïve and activated/memory Treg, we next addressed differences in activation potential of downstream pathways upon IL-6 treatment. Interestingly, STAT3 phosphorylation was induced in Treg with similar frequency and expression level in both naïve and activated Treg – even trending to higher pSTAT3 MFI in CD45RA^+^ Treg – despite different IL-6Rα expression levels **(Fig.3G**, **Fig. 4A-D)**. Treg were sorted as CD4^+^ CD25^+^ CD127^-^ cells with post-sort FOXP3 expression analysis **(Fig. 4E)** and were stimulated with IL-6 in expansion cultures containing IL-2 and CD3/anti-CD28 human T cell activation beads following protocols that are also used to expand human Treg *ex vivo* for clinical trial purposes^8^. IL-6 treated Treg did not show changes in viability or FOXP3 expression in both frequency and MFI on day 5 or day 8 of culture **(Fig. 4F-H)**, indicating that despite active signalling, Treg in expansion cultures maintained stable FOXP3 expression.

**Figure 4.**
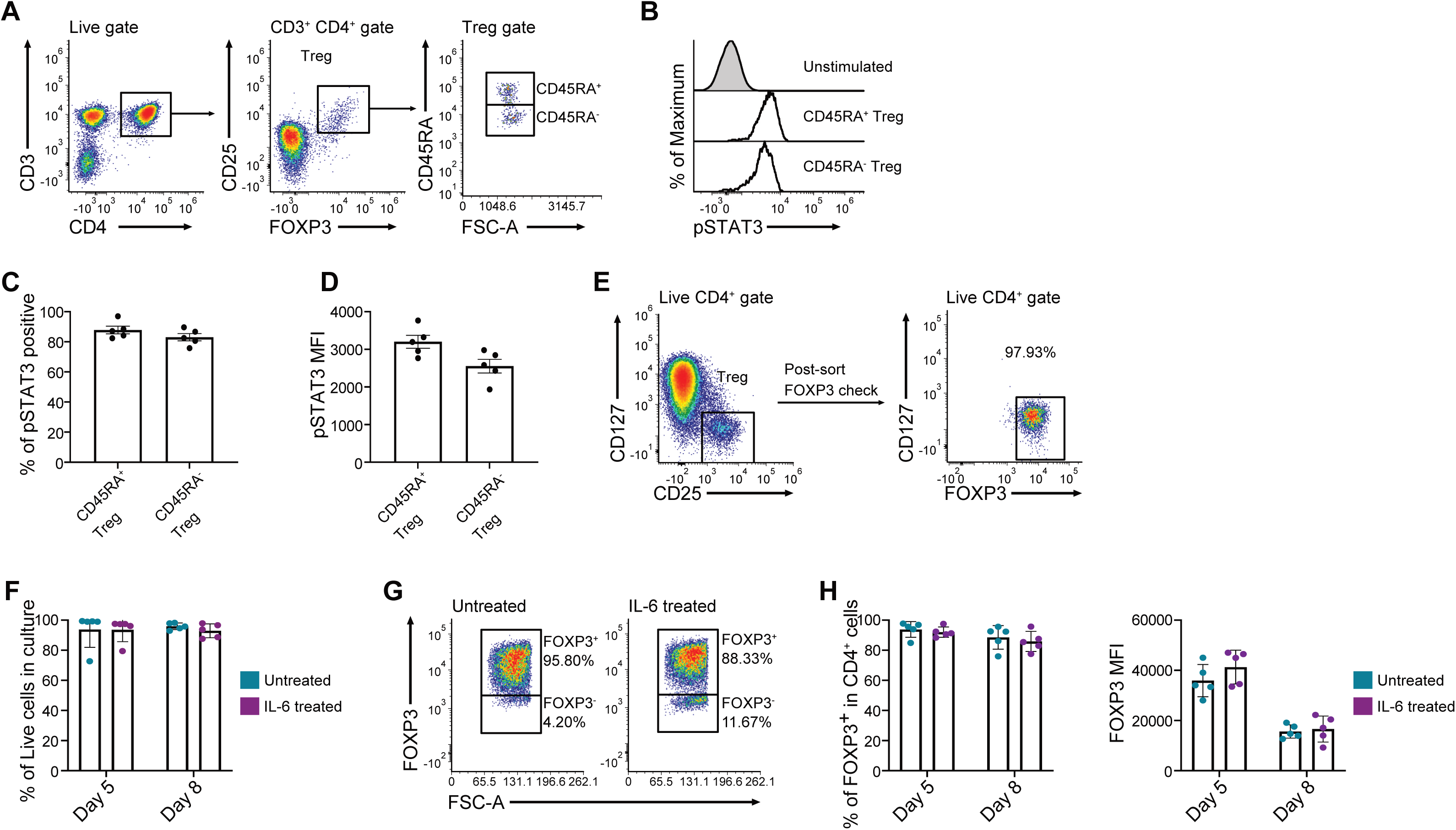
IL-6 can signal in both naïve and activated/memory Treg but does not affect FOXP3 stability. **(A)** Representative gating strategy to identify naïve and activated/memory Treg from PBMC stimulated with IL-6. **(B)** Histograms depicting pSTAT3 induction in naïve and activated/memory Treg upon IL-6 stimulation *in vitro*. Frequency **(C)** and MFI **(D)** of pSTAT3 induction Bar graph representations (mean ± SD, n=5) of frequencies as a percentage **(D)** and pSTAT3 MFI in naïve and activated/memory Treg upon IL-6 stimulation *in vitro* (mean ± SD, n=5). **(E)** Sort strategy to purify CD4+ CD25+ CD127-Treg from human PBMC, pre-gated for CD3+ live lymphocytes (left) and validation of FOXP3 expression in purified cells (right). **(F)** Frequency of live cells in Treg expansion cultures treated with or without IL-6 for 5 or 8 days (mean ± SD, n=5). FOXP3 expression levels, depicted as dot plots **(G)**, frequency or MFI **(H)**, in Treg expansion cultures treated with or without IL-6 for 5 or 8 days. [Statistical analysis was performed using Wilcoxon matched pairs signed rank test with FDR correction for multiple comparisons. Only significant comparisons are indicated.]

High concentrations of IL-2 and hence high levels of STAT5 activation may out-compete IL-driven STAT3 signalling^21^. We therefore tested whether IL-6 and pSTAT3 signalling could lead to FOXP3 instability in low IL-2 concentration conditions. Eight-day Treg expansion cultures were therefore cultured in varying concentrations of IL-2 (0 U/ml, 4 U/ml, 20 U/ml, 100 U/ml, 500 U/ml) and stimulated with either 0, 25 or 100 ng/ml of IL-6. As expected, total cell number and viability were strongly dependent on IL-2 concentrations. IL-6 stimulation also negatively affected total cell number and viability in a concentration dependent manner, although to a lesser extent than IL-2 starvation **(Fig 5A)**. However, the frequency of CD4^+^ FOXP3^+^ cells after 8 days of culture remained remarkably stable, both under limited IL-2 concentrations and in the simultaneous presence of high or intermediate levels of IL-6 **(Fig. 5B, C)**. These results indicate that while IL-6 treatment of Treg expansion cultures affected total cell number and viability, Treg maintained stable FOXP3 expression, even under limited IL-2 concentrations.

**Figure 5.**
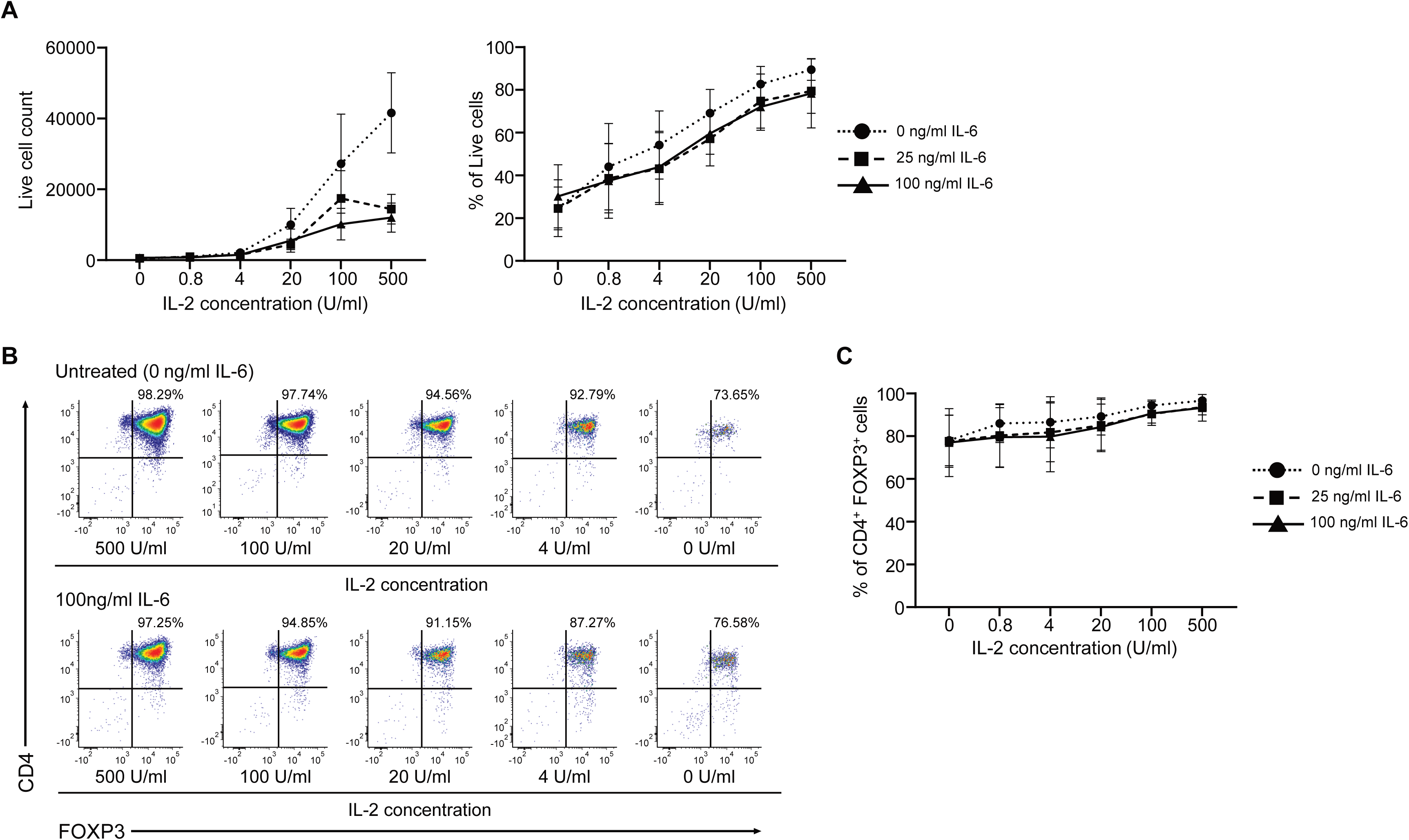
FOXP3 expression remains stable under limited IL-2 concentrations. **(A)** Line graphs of total viable cell number (left) and frequency of live cells (right) inTreg expansion cultures cultured with different IL-6 and IL-2 concentrations for 8 days (mean ± SD, n=4). Significant effects of IL-2 (p<0.0001) and IL-6 (p=0.0021) concentrations on the total number of viable cells and of IL-2 (p<0.0001) on the frequency of live cells were observed (Two-way ANOVA). **(B)** Representative flow cytometry plots of CD4 and FOXP3 expression levels in Treg expansion cultures cultured with different IL-6 and IL-2 concentrations for 8 days. **(C)** Line graph of CD4^+^ FOXP3^+^ cell frequency in Treg expansion cultures cultured with different IL-6 and IL-2 concentrations for 8 days (mean ± SD, n=4). Significant effects of IL-2 concentrations (p=0.0040) and IL-6 (p=0.0021) on CD4^+^ FOXP3^+^ frequencies were observed (Two-way ANOVA).

### FOXP3 expression remains stable upon stimulation with TGF-β and IL-6, but differentiates cultured Treg into Tr17

The functional adaptation Treg into specific T_H_-subset-suppressing cell types is an important hallmark of Treg biology^14–17^. In mice, IL-6 in conjunction with TGF-β has been shown to induce Tr17 from Treg, which have been implicated to play an important regulatory role in autoimmune diseases and to control pro-inflammatory T_H_17 responses^16,47–49^. Tr17 express Foxp3 alongside the T_H_17 lineage-defining marker RORγt and T_H_17-specific chemokine receptors. Given our findings that IL-6 did not induce FOXP3 instability, we assessed whether IL-6 signalling in conjunction with TGF-β could instead be used to differentiate Treg into Tr17. While IL-6 treatment alone did not induce RORγt expression in Treg expansion cultures (suggesting that the intrinsic production of TGF-β is not sufficient to drive Tr17 development), we observed a notable increase of RORγt expressing FOXP3^+^ cells in cultures treated with both TGF-β1 and IL-6 **(Fig. 6A, B)**. In line with the Treg lineage-stabilising nature of TGF-β1 signalling^50^, we also observed a concurrent increase of FOXP3 expression levels in these cultures compared to IL-6 treated or untreated conditions **(Fig. 6C)**. Of note, RORγt^+^ FOXP3^+^ Treg expressed higher levels of FOXP3 than their RORγt^-^ counterparts (which was not observed in CD4^+^ CD25^-^ CD127^+/-^ control cells), possibly implying a more stable and more suppressive phenotype^16,51,52^ **(Suppl. Fig. 2A, B)**. Similar to previous results **(Fig. 4H)**, IL-6 treatment did not lead to a decrease in FOXP3^+^ expression levels or the differentiation of FOXP3^-^ cells, further strengthening our conclusion that FOXP3 expression in human Treg is remarkably stable under IL-6/pSTAT3 signalling conditions.

**Figure 6.**
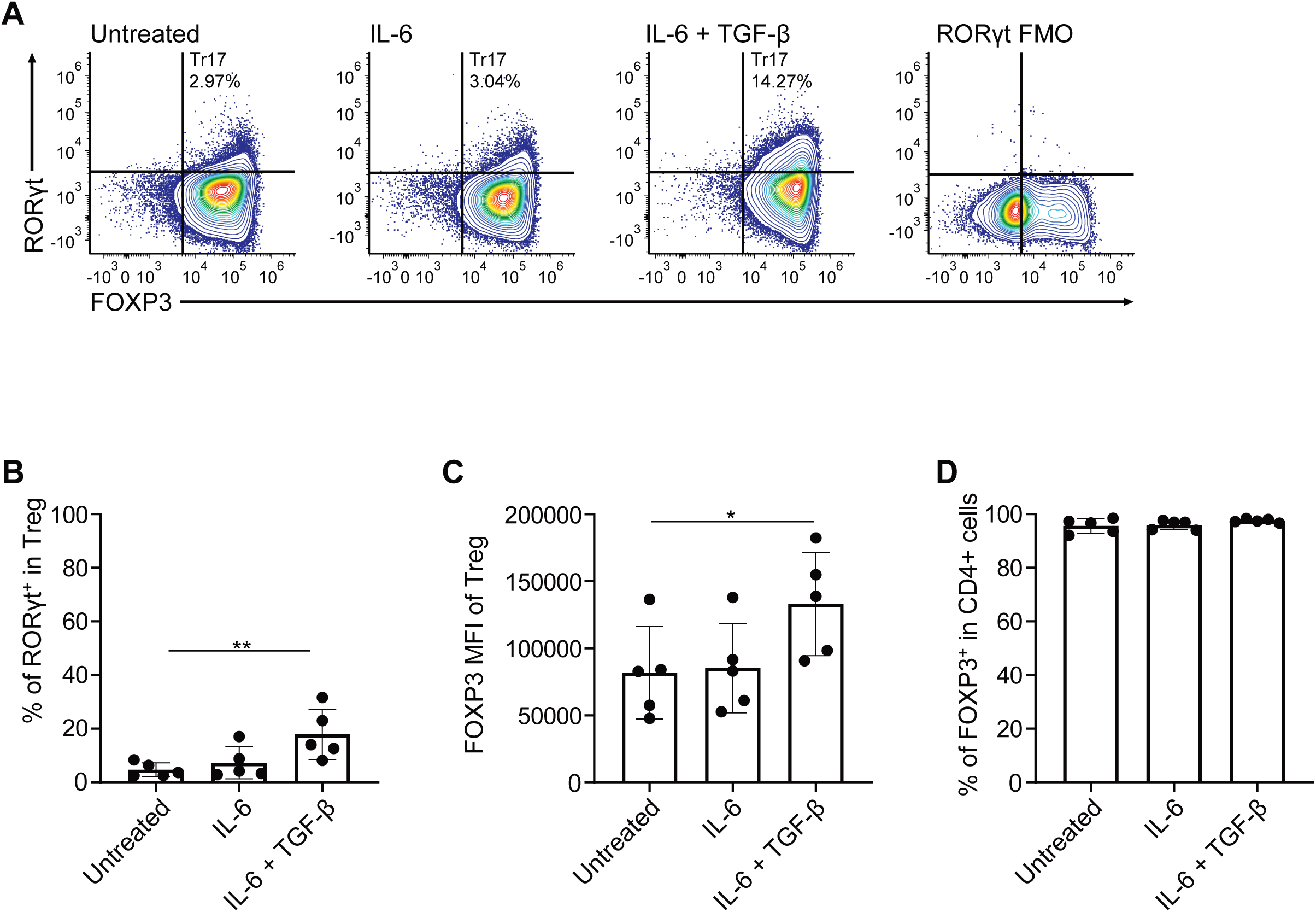
TGF-β and IL-6 induce Tr17 differentiation in Treg expansion cultures. **(A)** Representative FOXP3 and RORγT expression in Treg expansion cultures cultured under IL-6 (25 ng/ml) or TGF-β (10 ng/ml) and IL-6 (25 ng/ml) conditions for 6 days. **(B)** Frequency of RORγT^+^ expressing Treg **(B)**, FOXP3 expression levels in Treg **(C)** and total frequency of Treg **(D)** (mean ± SD, n=5) in Treg expansion cultures cultured under IL-6 (25 ng/ml) or TGF-β (10 ng/ml) and IL-6 (25 ng/ml) conditions for 6 days. [Statistical analysis was performed using Friedman test with post-hoc Dunn’s multiple comparisons. *p<0.05,**p<0.01; only significant comparisons are indicated.]

## Discussion

In this study, we examined the CD3^+^ CD4^+^ CD56^+^ CD16^+^ cells, a phenotype recently suggested to be acquired by exTreg in healthy human PBMC^30^. We demonstrate that the proposed exTreg signature, both at surface marker level and transcriptomic level, fails to differentiate these cells from NKT cells. Instead, the CD3^+^ CD4^+^ CD56^+^ CD16^+^ population is a diverse population mainly composed of cells expressing the NK(T) lineage marker PLZF and CD161 which can be further divided into naïve, effectory memory and central memory NKT cells. Additionally, we also observed a subpopulation of FOXP3^+^ CD25^+^ cells within the CD3^+^ CD4^+^ CD56^+^ CD16^+^ compartment, further outlining their heterogeneity and questioning their characteristic as exTreg. Similarly, the proposed ‘exTreg’^30^ and NKT cell^44^ transcriptomic signatures highly correlate with each other in CD3^+^ CD4^+^ CD56^+^ CD16^+^ PBMC from both healthy and inflammatory disease conditions.

We therefore postulate that rather than identifying exTreg, Freuchet et al. describe NKT in an atherosclerotic context. Several lines of evidence support this interpretation. CD4^+^ NKT cells have previously shown to play an inflammatory role in atherosclerotic Apoe^-/-^ mouse model by producing perforin and granzyme B^53^. It is possible that the Apoe^-/-^ FoxP3^eGFP−Cre-ERT2^ ROSA26^fl-stop-fl-tdTomato^ mouse model used by Freuchet et al. to detect exTreg, identified NKT that transiently upregulated Foxp3 due to the inflammatory microenvironment in atherosclerotic plaques, making them fate mapper/tdTomato-positive. The transient upregulation of FOXP3 has been shown in both human and murine iNKT, which can be mediated by CD3 activation or TGF-β signalling, respectively^37,38^. Although to our best knowledge Foxp3-expressing iNKT cells have not been studied in atherosclerotic mice, TGF-β has been shown to play a dominant role in driving murine atherosclerotic inflammation, with anti-TGF-β therapy actively suppressing lesion formation^54^.

While Freuchet et al. have carefully excluded the possibility that exTreg signatures describe NK cells, the same rigor has not been applied to NKT cells. Moreover, the TCRβ sequencing and GLIPH analysis performed by Freuchet et al. to analyse clonality^30^, does not incorporate NKT cell TCR signals, as GLIPH analysis considers MHC recognition for grouping TCR sequences^55^, thus neglecting CD1-d restricted NKT clonality as a comparative group.

Furthermore, we find no evidence of FOXP3 instability of human Treg in highly pro-inflammatory culture conditions, neither in the context of IL-6 (strongly associated with atherosclerosis^56^), TGF-β or under limiting IL-2 concentrations. Our data corroborate findings from Skartsis et al., demonstrating lineage stability of human Treg both *in vitro* and in humanized NSG mice upon treatment with IL-6 at high and low IL-2 concentrations^57^. Instead, we found that Treg expansion cultures treated with IL-6 in combination with TGF-β drove RORγt expression, while maintaining a continued high and robust FOXP3 expression. These results indicate that Treg cells can be differentiated into T_H_-suppressing phenotypes *in vitro*, and that these Tr cells might be utilized for an additional therapeutic benefit. Tr17 adoptive transfer therapy could be utilised to treat conditions like rheumatoid arthritis, which display a dominant T_H_-17 driven inflammatory profile^58^, similar to ongoing clinical trials focusing on T_H_2-like Treg adoptive cell therapy in ALS^59^.

Overall, our findings challenge the concept that Treg are unstable under pro-inflammatory conditions and rather indicate their robustness in Trge expansion cultures, especially in the context of IL-6-driven inflammation and polarisation to a Tr17 phenotype.

## Materials and Methods

### Isolation of human PBMC

Peripheral blood mononuclear cells (PBMC) were isolated from buffy coats of whole blood units of healthy donors using density gradient centrifugation. Blood was diluted with equal volume of cRPMI medium (RPMI Medium 1640, Gibco Cat No. 21875091; 10% FBS; 1% P/S) or undiluted, and was gently overlaid onto lymphocyte separation medium (LSM, MP Biomedicals, Cat No. 50494; or Biocoll, Bio&Sell, Cat No. BS.L 6115) at a 2:1 diluted blood to LSM ratio or 1:1 undiluted blood to Biocoll ratio, followed by centrifugation at 400 x g or 800 x g (no brake, minimum acceleration) for 20 to 30 minutes at 20°C. The PBMC interphase was aspirated, and the cells were washed with cRPMI or with wash solution (PBS + 2.5 mM EDTA) by centrifuging at 400 x g. RBC lysis was optionally performed by incubating the cells in 2ml of 1X RBC lysis buffer (RBC Lysis Buffer (10X); Biolegend, Cat No. 420302) for 10 minutes and a subsequent wash. For extended storage PBMC were transferred into cryovials (Cryo.S, greiner bio-one, Cat No. 122263) at a density of 30 x 10^6^ cells/ml in 1ml aliquots of freezing solution (10% DMSO, 90% FBS) and were frozen at a controlled rate using Mr. Frosty (Thermo Scientific, Cat No. 51000001) at -80°C for 24 hours. The PBMC cryovials were subsequently transferred for extended storage in liquid nitrogen.

### Treg sorting from human PBMC

Cryopreserved PBMC were thawed at 37 °C and resuspended in FACS buffer (1x PBS from 10X DPBS, Gibco, Cat No. 1420067, 5% FBS) at 4 °C. PBMC samples were centrifuged at 100 x g for 15 minutes and the pellet was resuspended in FACS buffer three times for platelet depletion. The thawed cells were allowed to recover protein expression before Treg sorting by placing them in a high-density culture (HDC). For HDC, the samples were resuspended in cRPMI at a density of 1 x 10^7^ cells/ml incubated at 37 °C, 5% CO_2_ for 24 to 48 hours. After resting the PBMC in HDC, CD4 cells were enriched using the MojoSort Human CD4 T cell isolation kit (Biolegend, Cat No. 480130) by following the manufacturer’s protocol. 1:4 of the manufacturer’s recommended volumes for Biotin-Antibody cocktail and Streptavidin Nanobeads were used.

Enriched CD4 cells were subsequently depleted of CD127^+^ cells by staining with anti-human CD127-PE antibody (clone A019D5, Biolegend) and using the MojoSort Human anti-PE nanobeads (Biolegend, Cat No. 480092) following the manufacturer’s protocol. MojoSort buffer recommended by the manufacturer was substituted with FACS buffer and all the centrifugation steps were performed at 400 x g and 4 °C. For Treg sorting, the cells were stained with anti-human CD4-APC (clone RPA-T4, Biolegend), anti-human CD127-PE (clone A019D5, Biolegend), and co-stained with three anti-human CD25-BV711 antibodies (clone 2A3, BD Biosciences; clone BC96, Biolegend; clone MA251, Biolegend). Treg were sorted as CD4^+^ CD25^+^ CD127^-^ cells using the Sony MA900 sorter. Post-sort purity was verified by re-analysis of the sorted cells and Treg purity was verified by FOXP3 staining.

### *In vitro* culture of sorted human Treg

Sorted Treg were cultured in cRPMI supplemented with rIL-2 (Proleukin, Clinigen) at a concentration of 500 U/ml unless otherwise indicated and activated by anti-CD3/anti-CD28 beads (Dynabeads Human T activator CD3/CD28, Gibco, Cat No. 11161D) at 1:1 cell to bead ratio. The cells were seeded at a maximum density of 1 x 10^5^ cells per well in 200 µl of Treg culture media in a 96-well U-bottom cell culture plate (Greiner Bio-one, Cat No. 650180) and incubated at 37 °C, 5% CO_2_. The culture wells with T_H_-polarizing/inflammatory test conditions were additionally supplemented with recombinant human IL-6 (Biolegend, Cat No. 570802; Miltenyi, Cat No. 130-095-365) or combination of human IL-6 and human TGF-β1 (Miltenyi, Cat No. 130-095-067) at indicated concentrations. The media was refreshed every two days and half of the media was replaced with fresh media on the other days.

### IL-6 stimulation assay and pSTAT3 staining

Cryopreserved PBMC were thawed and placed in HDC for 24 to 48 hours. The cells were incubated with fixable viability dye (LiveDead Fixable Blue, Invitrogen, L23105), and Human Fc block (BD Pharmingen, Cat No. 564220) for 15 minutes in 1X PBS, and stained with anti-human CD4-BUV496 (clone: SK3), and anti-human CD25-BV711 (clone: BC96) antibodies in FACS buffer for 45 minutes. The cells were resuspended in cRPMI without FBS and plated in 96-well U-bottom culture plates with each well containing 5 x 10^5^ cells in 200ul and incubated at 37 °C, 5% CO_2_ for 3 hours. After the incubation period, the cells were centrifuged and resuspended in fresh cRPMI without FBS in 96-well V-bottom plates (Thermo Scientific, Cat No. 611V96) and 50 ng/ml of IL-6 (Miltenyi, Cat No. 130-095-365) was added to the wells and incubated at 37 °C, 5% CO_2_ for 30 minutes. The cells were fixed by adding equal volume of 4% paraformaldehyde and incubating for 20 minutes. The cells were subsequently washed with FACS buffer and permeabilized in ice-cold MeOH for 20 minutes and washed using FACS buffer. The fixed and permeabilized cells were stained overnight with anti-pSTAT3-Pacific Blue (4/P-STAT3, BD, Cat No. 560312), anti-human FOXP3, anti-human CD3, anti-human CD31-PE/Dazzle 594, and anti-human CD45RA antibodies (refer antibody table).

### Flow cytometry

Antibody panels were developed using established panel design strategies optimised for spectral flow cytometry on the Cytek Aurora^60^. Cells to be stained with the antibody panels were plated in 96-well V-bottom plates and incubated with a fixable viability dye (Zombie NIR or LiveDead Fixable Blue) and FcR block (Human Trustain FcX, Biolegend, Cat No. 422301; or BD Pharmingen Human Fc block) diluted in 1x PBS for 15 minutes and were stained with the surface antibody panel diluted in FACS buffer and incubated for 45 minutes. The cells were subsequently washed twice, and the intracellular staining was performed overnight using the eBioscience Foxp3/Transcription factor buffer set (Invitrogen, Cat. No. 00552300) by following the manufacturer’s protocol. The stained samples were acquired on the Cytek Aurora (4 laser 16UV-16V-14B-8R or 3 laser 16V-14B-8R), and the fcs files were analysed using FCS express (version 7.16.0035). The antibodies used are listed in Table 1.

**Table 1.** Flow cytometry staining panel. List of antibodies and concentrations used for surface and intracellular flow cytometry staining. Details of antibodies used for pSTAT3 staining and Treg sorting are provided in the respective methods sections.

| Panel | Antibody | Fluorochrome | Clone | Vendor | Final conc. |
| --- | --- | --- | --- | --- | --- |
| Surface<br>4°C<br>staining | Anti-hu TIGIT | BV480 | 741182 | BD Biosciences | 4 µg/ml |
|  | Anti-hu CD25 | BV711 | BC96 | Biolegend | 2 µg/ml |
|  | Anti-hu CD39 | BUV563 | TU66 | BD Biosciences | 4 µg/ml |
|  | Anti-hu LAG3 | BV421 | 11C3C65 | Biolegend | 5.6 µg/ml |
|  | Anti-hu GITR | APC-Fire 750 | 108-17 | Sony | 8 µg/ml |
|  | Anti-hu CD127 | PE-Cy7 | eBioRDR5 | Sony | 1 µg/ml |
|  | Anti-hu CCR4 | PE-Fire 700 | L291H4 | Sony | 1 µg/ml |
|  | Anti-hu CD56 | BV750 | 5.1H11 | Sony | 0.25 µg/ml |
|  | Anti-hu CD27 | Vio Bright R720 | REA499 | Miltenyi Biotec | 5 µg/ml |
|  | Anti-hu CD31 | PE | WM59 | Biolegend | 1 µg/ml |
|  | Anti-hu CD31 | PE-Dazzle 594 | WM59 | Biolegend | 1 µg/ml |
|  | Anti-hu CD45RA | PerCP-eFluor710 | GRT22 | eBioscience | 0.96 µg/ml |
|  | Anti-hu CD294 | BV421 | BM16 | Biolegend | 1.5 µg/ml |
|  | Anti-hu CD16 | NovaFluor Blue 585 | 3G8 | eBioscience | 1.2 µg/ml |
|  | Anti-hu CD25 | BV480 | M-A251 | BD Biosciences | 2 µg/ml |
|  | Anti-hu CD45RO | BV750 | UCHL1 | Biolegend | 0.12 µg/ml |
|  | Anti-hu CD56 | PE-Fire 640 | QA17A16 | Biolegend | 0.25 µg/ml |
|  | Anti-hu TIGIT | PE-Fire 810 | A15153G | Biolegend | 1 µg/ml |
|  | Anti-hu CD127 | Spark NIR 685 | A019D5 | Biolegend | 1 µg/ml |
|  | Anti-hu PD-1 | APC-Fire 810 | A17188B | Biolegend | 0.5 µg/ml |
|  | Anti-hu CD19 | Biotin | HIB19 | Biolegend | 5 µg/ml |
|  | Anti-hu CD14 | Biotin | M5E2 | Biolegend | 5 µg/ml |
|  | Anti-hu CCR7 | APC-Cy7 | G043H7 | Biolegend | 1 µg/ml |
|  | Dump: streptavidin | BV650 | - | Biolegend | 0.25 µg/ml |
|  | Anti-hu CD126 | PE | UV4 | Biolegend | 8 µg/ml |
| Intracellular staining | Anti-hu CTLA-4 | BV605 | BNI3 | Sony | 0.5 µg/ml |
|  | Anti-hu CD3 | Spark Blue 550 | SK7 | Sony | 0.25 µg/ml |
|  | Anti-hu FOXP3 | Alexa Fluor 488 | 206D | Sony | 0.5 µg/ml |
|  | Anti-hu CD4 | eFluor 450 | OKT4 | eBioscience | 0.125 µg/ml |
|  | Anti-hu CD8 | PerCP | SK1 | Sony | 0.25 µg/ml |
|  | Anti-hu RORgt | PE | Q21-559 | BD Biosciences | 0.25 µg/ml |
|  | Anti-hu CD161 | Pacific blue | HP-3G10 | Biolegend | 0.5 µg/ml |
|  | Anti-hu CD3 | BV510 | SK7 | Biolegend | 0.4 µg/ml |
|  | Anti-hu CD27 | BV570 | O323 | Biolegend | 0.5 µg/ml |
|  | Anti-hu Ki-67 | BV605 | Ki-67 | Biolegend | 0.0625 µg/ml |
|  | Anti-hu CD45 | BV785 | HI30 | Biolegend | 0.0625 µg/ml |
|  | Anti-hu TIM-3 | FITC | F38-2E2 | Biolegend | 1 µg/ml |
|  | Anti-hu PLZF | PE | R17-809 | BD Biosciences | 1 µg/ml |
|  | Anti-hu FOXP3 | PE-Dazzle 594 | 206D | Biolegend | 0.8 µg/ml |
|  | Anti-hu CD103 | PerCP-eFluor710 | B-Ly7 | eBioscience | 0.0625 µg/ml |
|  | Anti-hu Helios | PE-Cy7 | 22F6 | Biolegend | 0.015 µg/ml |
|  | Anti-hu CD4 | BUV737 | SK3 | BD Biosciences | 1:3200 dilution |
|  | Anti-hu CD8 | AF700 | SK1 | Biolegend | 0.0156 µg/ml |

### Flow cytometry data analysis

Flow cytometry data was unmixed in the SpectroFlo software, and the unmixed fcs files were analysed in FCS Express (version 7.16.0035). For multiparametric analysis using FlowSOM, cells of interest were manually gated and exported into R (version 4.4.3) using RStudio (version 2024.12.0+467). Subsequently, the markers for clustering were indicated and arcsinh transformation was performed using the flowVS package. The cell counts of donors for analysis were not normalised in CD4^+^ CD56^+^ CD16^+^ cell phenotyping and cell counts of donors for CD4^+^ FOXP3^+^ CD25^+^ Treg were normalised to the sample with the least number of Treg. FlowSOM clustering was performed on the transformed files using the CATALYST package^61,62^.

### CITE-seq data analysis

Analysis was performed on five publicly available PBMC CITE-seq datasets: Coronary artery disease (CAD; GSE190570), Healthy volunteers in HIV-vaccine trial (VAC vaccine trial; GSE164378), Type 1 diabetes (T1D; GSE201197), Psoriatic Arthritis (PsA; GSE280319), Immune dysregulation due to ARPC5 mutation (ARPC5mut; GSE215451).

For each dataset, pre-processed RNA expression matrices, antibody-derived tag (ADT) counts and associated metadata were obtained and imported into R for downstream analysis using the seurat package (version 5.3.0). ADT data were normalised using the centered log-ratio (CLR) transformation. CD3^+^ CD4^+^ CD56^+^ CD16^+^ CD8^-^ CD19^-^ CD14^-^ cell subset was selected based on ADT expression profiles. RNA counts were normalised using the seurat’s NormalizeData() function. Dimensionality reduction was performed using principal component analysis (PCA), followed by Uniform Manifold Approximation and Projection (UMAP). Graph-based clustering was performed using the shared nearest neighbor (SNN) modularity optimization algorithm implemented in Seurat (FindClusters()).

### Statistical analysis

Statistical analysis was performed in GraphPad Prism (version 9.5.1). The Wilcoxon matched pairs signed rank test was performed to compare two paired groups and multiple Wilcoxon matched pairs signed rank tests was used to compare two paired groups within different treatment conditions. Friedmann test with post-hoc Dunn’s multiple comparison test was performed to compare more than two paired groups. For analysing the effects of two factors and their interaction within paired samples, a two-way repeated measures ANOVA was performed, followed by post-hoc Tukey’s multiple comparisons test. A p-value of 0.05 was considered significant.

## Acknowledgements

We greatly appreciate the contributions by Adamantios Mavrogiannis^†^ from the Adaptive Immunology Laboratory, KU Leuven. The authors further acknowledge the contributions of the KUL Flow and Mass Cytometry core facility for flow cytometry and cell sorting support and the services of the Transfusion Center of the University Medical Center Mainz, Johannes Gutenberg University, Mainz, Germany. This work was funded by the Research Foundation Flanders (FWO) (Fundamental Research Grant, G054722N) to S.M.S. and supported by the Rise up! programme of the Boehringer Ingelheim Foundation (BIS) and the German Research Foundation (DFG) Research Units Programme (FOR) 5644, project no. 515636567 to J.U.M.

## Author contributions

S.M.S. and J.U.M. acquired funding, conceived, designed and supervised the study, interpreted data, and wrote the manuscript. J.M. performed experiments, analysed and interpreted data, and wrote the manuscript. M.K. performed experiments, analysed and interpreted data. M.S. performed bioinformatic analysis. All authors commented on and approved the final manuscript.

---

**Supplementary figure 1.**
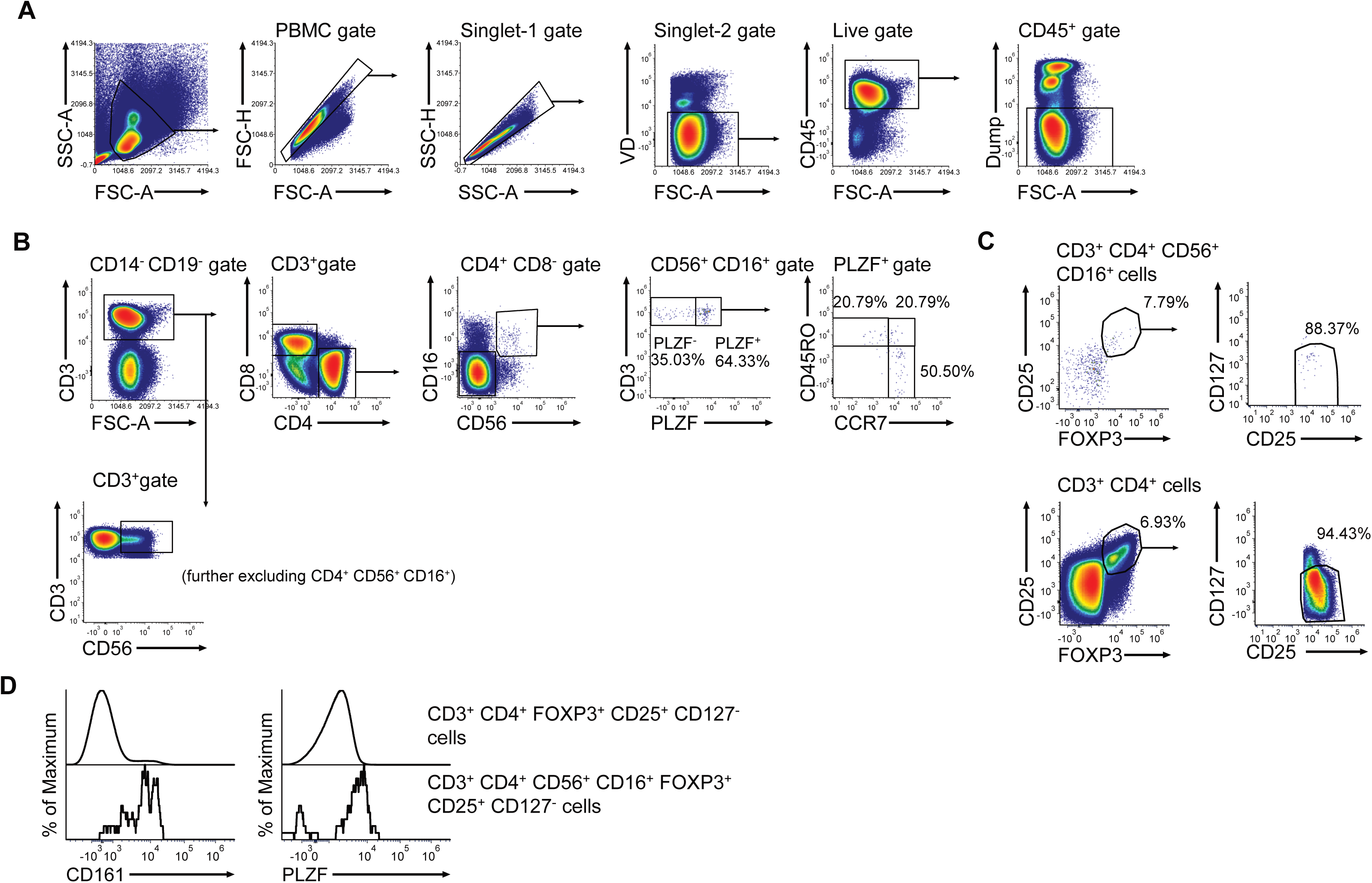
Gating strategy for CD4+ CD56+ CD16+ phenotyping panel and Treg marker expression. **(A)** Representative flow cytometry graph of pre-gating strategy before gating target populations. **(B)** Gating strategy for analysing CD4+ CD56- CD16- population, CD4+ CD56+ CD16+ population and CD3+ CD56+ population (excluded CD4+ CD56+ CD16+, not shown). CD4+ CD56+ CD16+ population was analysed based on PLZF expression, followed by CD45RO and CCR7 expression. **(C)** Flow cytometry gating for Treg markers FOXP3+, CD25+ and CD127-expression in CD3+ CD4+ CD56+ CD16+ population and CD3+ CD4+ population. Data from 5 PBMC donors were merged and presented as a single dataset. **(D)** Histograms comparison of CD161 and PLZF expression in CD3^+^ CD4^+^ FOXP3^+^ CD25^+^ CD127^-^ and CD3^+^ CD4^+^ CD56^+^ CD16^+^ FOXP3^+^ CD25^+^ CD127^-^ populations. Data from 5 PBMC donors were merged and presented as a single dataset.

**Supplementary figure 2.**
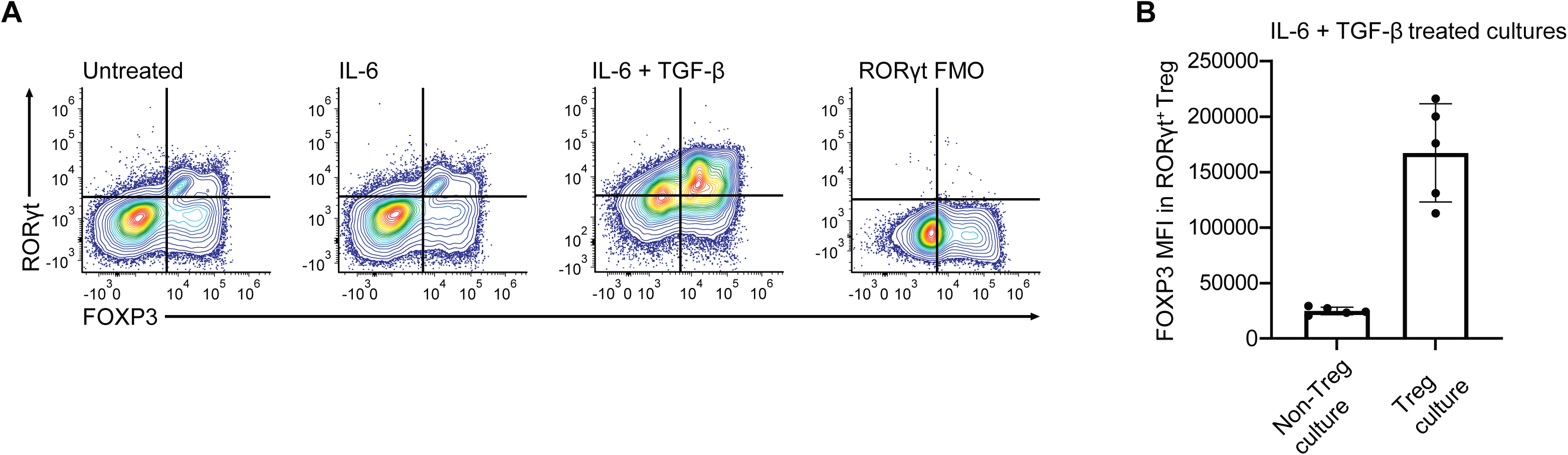
Non-Treg expansion culture with TGF-β and IL-6. **(A)** Representative flow cytometry graph of FOXP3 and RORγT expression upon treatment and expansion of sort-purified CD4^+^ CD25^-^ CD127^+^ non-Treg with 10 ng/ml TGF-β + 25 ng/ml IL-6, IL-6 and untreated, respectively. **(B)** Bar graph representation (mean ± SD, n=5) of FOXP3 MFI in RORγT^+^ Treg (FOXP3^+^) in Treg and non-Treg cultures stimulated with 10 ng/ml TGF-β + 25 ng/ml IL-6. Wilcoxon matched pairs signed rank test was performed to compare FOXP3 MFI of RORγT^+^ Treg in Non-Treg and Treg cultures. The experiment was performed in duplicates, and the data is represented as the mean of measurements (frequency, MFI) of the duplicates.

